# A benchmark comparison of CRISPRn guide-RNA design algorithms and generation of small single and dual-targeting libraries to boost screening efficiency

**DOI:** 10.1101/2024.05.17.594311

**Authors:** Sebastian Lukasiak, Alex Kalinka, Nikhil Gupta, Angelos Papadopoulos, Khalid Saeed, Ultan McDermott, Gregory J Hannon, Douglas Ross-Thriepland, David Walter

**Affiliations:** Joint AstraZeneca-Cancer Research Horizons Functional Genomics Centre, Cambridge, UK; Discovery Sciences, BioPharmaceuticals R&D, AstraZeneca, Cambridge, UK; Cancer Research Horizons, Cambridge, UK; Oncology R&D, AstraZeneca, Cambridge, UK; Cancer Research UK, Cambridge Institute, Cambridge, UK

**Keywords:** Pooled CRISPR screening, Genome-wide, CRISPR-Cas9, sgRNA selection algorithm

## Abstract

Genome-wide CRISPR sgRNA libraries have emerged as transformative tools to systematically probe gene function. While these libraries have been iterated over time to be more efficient, their large size limits their use in some applications. Here, we benchmarked publicly available genome-wide single-targeting sgRNA libraries and evaluated dual targeting as a strategy for pooled CRISPR loss-of-function screens. We leveraged this data to design two minimal genome-wide human CRISPR-Cas9 libraries that are 50% smaller than other libraries and that preserve specificity and sensitivity, thus enabling broader deployment at scale.

## Background

CRISPR-based loss-of-function pooled screens have revolutionized our ability to systematically interrogate gene function. The sensitivity of these screens depends critically on the efficiency with which sgRNAs create loss-of-function alleles. Genome-wide CRISPR-Cas9 sgRNA libraries have been iteratively optimised to reduce off-target activity and increase on-target efficiency (1-7). Dual targeting libraries, where two sgRNAs are used per gene, were suggested to perform better than single targeting libraries (8, 9). However, there is no clear consensus on which library performs best for pooled loss-of-function screening and how dual targeting libraries perform when compared to conventional single targeting libraries. Applying large libraries that contain many constructs per gene is cost intensive and not feasible for some applications (e.g., organoids, in vivo). Smaller genome-wide CRISPR libraries are more cost-effective and increase feasibility when assaying complex models. We hypothesised that libraries with fewer constructs per gene that are chosen according to principled criteria and effectively modulate targeted genes perform as well or better than larger libraries. To test this, we systematically evaluated various sgRNA libraries, including an assessment of dual targeting. We leveraged these insights to create two minimal genome-wide CRISPR libraries and conducted concordance screens to assess their performance.

## Results and Discussion

The overlap of sgRNAs between commonly used CRISPR libraries is relatively small (Figure S1A, B) and it is not clear which library performs best in CRISPR-based loss-of-function screens. To fairly compare the performance of libraries we assembled a benchmark human CRISPR-Cas9 library comprised of gRNA sequences targeting 101 early essentials, 69 mid essentials, 77 late essentials (10) and 493 non-essentials (see Methods; Figure S1F). The gRNA sequences were taken from 6 pre-existing libraries i.e. Brunello (1), Croatan (8), Gattinara (11), Gecko V2 (5), Toronto v3 (4), and Yusa v3 (12), (Figure S1). Comparison of their sequences indicates that each library has a higher proportion of private as opposed to shared guides (Figure S1B). Essentiality screens were performed in HCT116, HT-29, RKO, and SW480 colorectal cancer cell lines (see Methods, Figure 1A). As sgRNA Vienna Bioactivity CRISPR (VBC) scores have been calculated genome-wide for all coding sequences (13), we used these scores to identify the top 3 (henceforth referred to as ‘‘top3-VBC’’) and bottom 3 guides (henceforth referred to as ‘‘bottom3-VBC’’) in our benchmark library. The results of the pooled CRISPR lethality screens performed in the four cell lines showed that the top3-VBC guides exhibit the strongest depletion curves while the bottom3-VBC guides have the weakest depletion curves, with all other libraries sitting in-between these two bounds, with Yusa and Croatan as the best performing libraries (Figure 1A, B). Similar results were observed across early, mid, and late essentials (Figures S2, S3).

**Figure 1.**
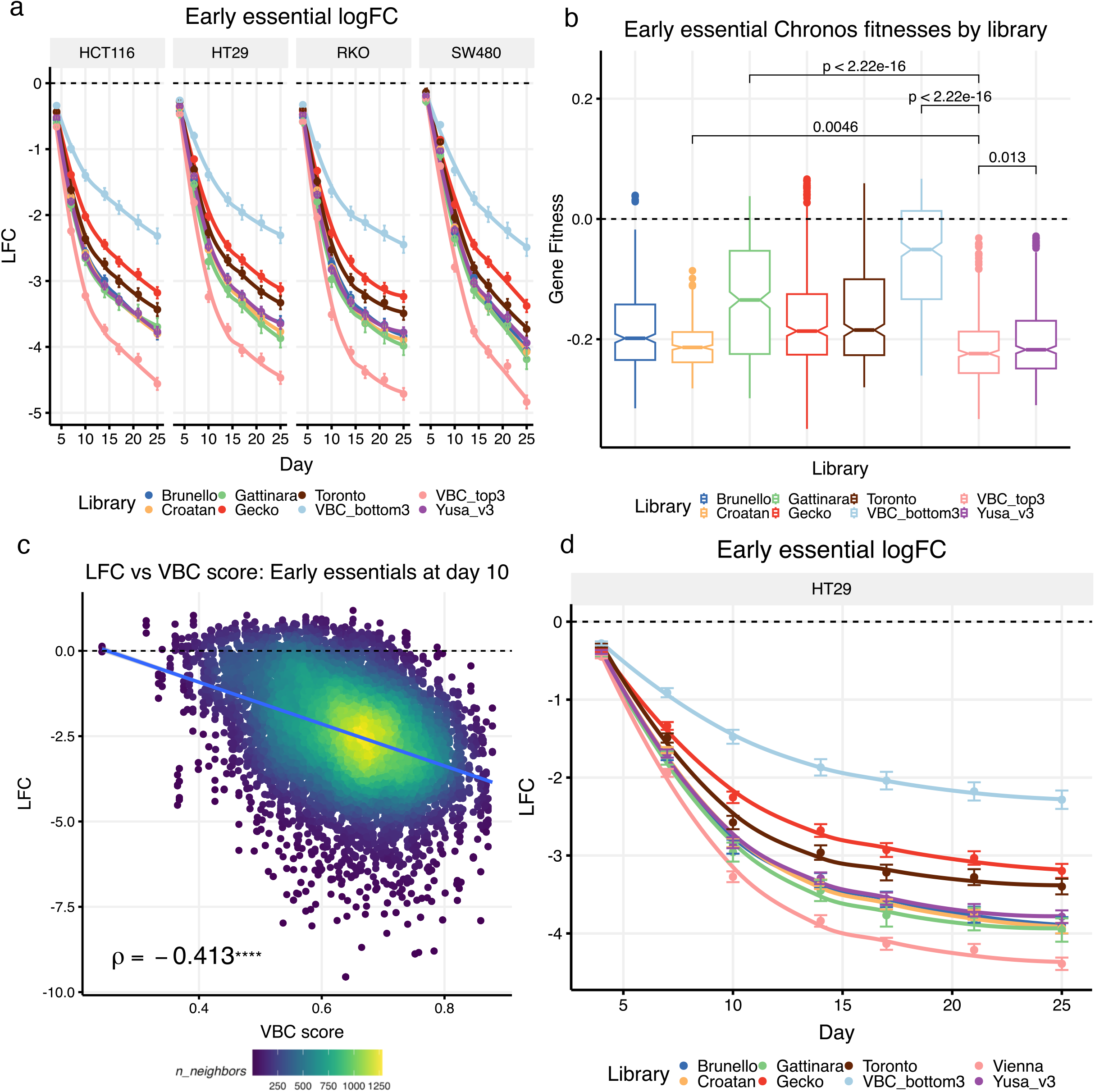
A three-guide-per-gene CRISPRn library performs as well or better than libraries containing more guides per gene. a) Early essential gene depletion curves at the gRNA level across four cell lines. The results show that the three guides with the highest VBC scores (‘VBC_top3’) among the six main libraries (Brunello, Croatan, Gattinara, Gecko, Toronto, and Yusa v3) exhibit the strongest depletion, and the three guides with the lowest VBC scores (‘VBC_bottom3’) have the weakest depletion. Points indicate averages across gRNAs (+/-standard error of the mean). LOESS curves are fitted to the data. b) Chronos gene fitness estimates for early essential genes pooled across the four cell lines shown in panel ‘a’. P-values for selected pairwise comparisons between ‘VBC_top3’ and four other libraries are shown (Wilcoxon two-sample tests). c) Early essential correlation between log_2_ fold-change and VBC scores (at day 10 across all four cell lines in panel ‘a’) shows a significant negative correlation (Spearman’s rho shown). d) Early essential gene depletion in a benchmark library containing the top-6 VBC guides (the ‘Vienna’ library) reproduces the results shown in panel ‘a’ in a single cell line. LFC -log_2_ fold-change.

The Chronos algorithm models CRISPR screen data as a time series producing a single gene fitness estimate across all the time points sampled in the experiment (14). Gene fitness estimates measured this way show that the 3-guides-per-gene of the top3-VBC library is no worse than the two best performing libraries with more guides per gene – Yusa (average of 6 guides per gene), Croatan (average of 10 guides per gene). The bottom3-VBC is once again the worst performing library (Figures 1B, S4). Consistent with this result, VBC scores correlate negatively with the log-fold changes of guides targeting essential genes and therefore provide a means to predict gRNA efficacy, albeit not perfectly (Figure 1C). It is worth noting that the recently developed Rule Set 3 scores (7) also exhibit a negative correlation with log fold changes (Figure S5A), and these two scores also correlate with one another (Figure S5B).

In a follow-up study, we modified the original benchmark library to include the top 6 VBC gRNAs per gene (henceforth referred to as the ‘Vienna’ library) and repeated the lethality screen in HT-29 cell line (Figure 1D). The results show that the Vienna library has the strongest depletion curve (Figures 1D, S6, S7). We also identified the guides in our benchmark library that overlapped with guides in the MiniLib-Cas9 (MinLib) 2-guide library (3) and found that the MinLib library is the best performer in terms of essential gene depletion (Figure S8).

To compare single- and dual-targeting strategies, we created another benchmark library, ‘‘benchmark-dual human CRISPR-Cas9 library’’ using the same genes and guides used in the first ‘‘benchmark human CRISPR-Cas9 library’’ but paired so that both guides in each guide pair target the same gene (see Methods). Guides in this library were also paired with Non-Targeting Controls (NTCs) so that we could directly compare single- and dual-targeting guide pairs in the same screen (Figures S9, S10, S11). We used this library to conduct a lethality screen in HCT116, HT-29 and A549 cell lines and the results showed that depletion of essential genes was on average strongest in the dual-targeting guide pairs relative to the single-targeting pairs (Figure 2A). Such a benefit of dual targeting was previously reported (8) and attributed to a deletion between the two sgRNA target sites creating a knockout more effectively than error prone repair in response to a single sgRNAs mediated DNA double strand break. However, as well as stronger depletion of essentials, the dual-targeting guides also exhibited on average weaker enrichment of non-essential genes relative to the single-targeting guides (as measured by log-fold changes – Figure S10B; Chronos gene fitness estimates also indicate weaker enrichment of non-essentials in the dual-targeting guides – Figure 2B). It is unlikely that there is a fitness cost associated with targeting these genes as they are classified as neutral (15) and have zero expression in CCLE data in the relevant cell lines (Figure S12). However, for these neutral genes we estimated that there is, on average, a log_2_-fold change delta of –0.9 (dual minus single) that is relatively constant across time points (Figure S12). This could reflect a dual-targeting fitness cost associated with creating twice the number of dsDNA cuts in the genome and highlights that some caution may be warranted when electing to use a dual-targeting library as the potential triggering of a heightened DNA damage response may be undesirable in certain CRISPR screen contexts. We further observed that the essential-gene depletion advantage enjoyed by the Vienna single-targeting library was largely ablated in the dual-targeting screen (Figure S9) implying that the benefit of dual-targeting may be greater when the knock-out performance of less efficient guides is compensated by pairing them with more efficient guides. We further note the absence of a clear impact of the distance between gRNA pairs, measured either in absolute terms or relative to gene length (Figure S13) as was previously reported (16, 17). To test the performance of smaller genome-wide libraries designed using VBC scores, we performed a genome-wide Osimertinib drug-gene interaction resistance screen using a library composed of the top 3 VBC guides per gene (henceforth referred to as Vienna-single) and the Yusa v3 6-guide library (Yusa) in HCC827 and PC9 lung adenocarcinoma cell lines (see methods). We also ran this screen using a dual-targeting library in which the top 6 VBC guides were paired in specific combinations such that both guides in each guide pair target the same gene (henceforth referred to as Vienna-dual) (see methods). The lethality results in the control arm both confirm and extend the lethality results found in the benchmark screen with the Yusa 6-guide library consistently performing the worst (Figures 2C, 2D).

**Figure 2.**
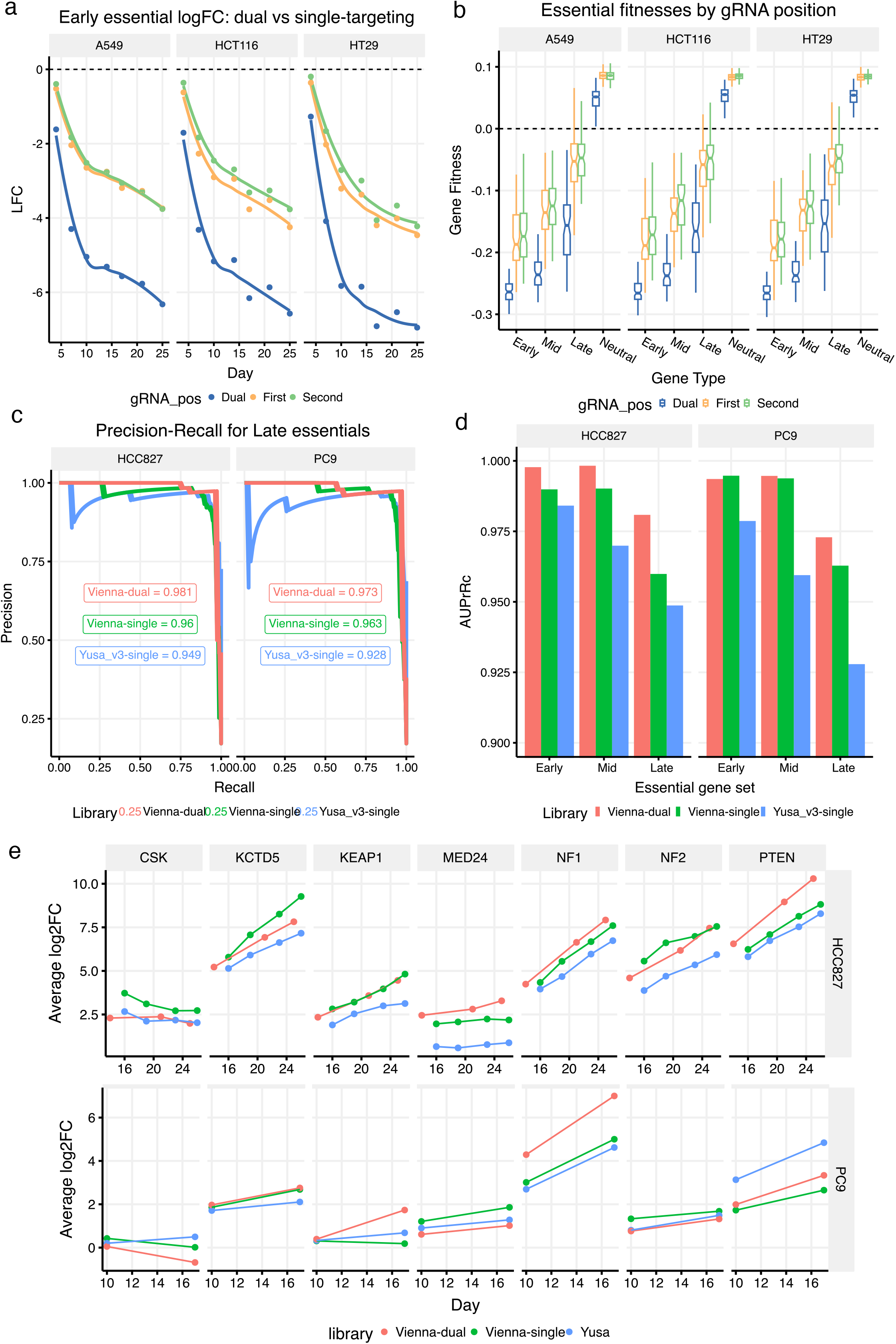
Small dual-guide CRISPRn libraries perform as well or better than their single-guide equivalents. a) Early essential gene depletion curves at the gRNA level for a dual-guide benchmark library across three cell lines. The results show that dual-targeting guides (‘Dual’) out-perform single-targeting guides when they are paired with NTCs in either the ‘First’ or ‘Second’ position. Points indicate averages across gRNAs (+/-standard error of the mean). LOESS curves are fitted to the data. ‘gRNA_pos’ -gRNA position. b) Chronos gene fitness estimates for the dual-guide benchmark library showing that the dual-targeting guides out-perform the single-targeting guides across all essential gene groups. For neutral genes, however, dual-targeting guides exhibit weaker fitness estimates than the single-targeting guides. c) Precision-Recall curves for Chronos fitnesses of late essential genes in three separate genome-wide screens across two cell lines. The area under the precision recall curves is shown for each library. d) Area under the precision recall curves for early, mid, and late essential gene sets broken down by library and cell line. AuPrRc – area under precision recall curve. e) Log_2_ fold-change dynamics of treatment samples relative to the plasmid in three separate genome-wide Osimertinib screens for a panel of seven genes that were independently validated as resistance hits. The results show that dual-guide and single-guide libraries with three guide constructs per gene perform comparably to the six-guide Yusa v3 library. Results shown for two cell lines.

Improved performance in terms of depletion of essentials also carries over to improved performance in the drug-gene interaction comparisons (Figures 2E). A fair assessment of performance was afforded by the existence of seven independently validated resistance hits from the original EGFR screen (19). In both cell lines, the Vienna-single and Vienna-dual libraries exhibited the strongest resistance log fold changes for the validated resistance genes with Yusa the strongest in only one case out of 14 in total and being consistently the lowest in 9 of the remaining 13 (Figures 2E). Taking 100 resistance hits called by either MAGeCK (20) or a Chronos two-sample analysis ((14); see Methods) and ranking them by either their log-fold changes or their Chronos gene fitness delta (treatment minus control), we found that the Vienna-dual library consistently exhibited the highest effect size across both cell lines (Figures S14, S15).

## Conclusions

This study demonstrates that CRISPR screen performance of both lethality and drug-gene interaction screens can be as good or better in smaller libraries when the guides are chosen according to principled criteria. The major consequence of this finding is that screens conducted with small libraries could be run at a smaller scale without compromising performance and this would entail a concomitant reduction in cost and an increase in throughput. We further demonstrate performance enhancements in dual-targeting libraries, both for lethality and drug-gene interaction screens, but caution that the use of such libraries may not always be suitable in the context of specific CRISPR screens.

## Methods

### Library designs

To select essential genes for the benchmarking libraries ‘‘benchmark human CRISPR-Cas9 library’’ and ‘‘benchmark-dual human CRISPR-Cas9 library’’, we used two criteria: 1) membership in the Sanger pan-essential gene list (21) and/or Hart essential gene list (15), and 2) classification as early, mid, or late essential based on (10). Neutral genes were taken from the Hart list (15). After selecting genes represented in all component libraries, the final numbers were: 101 early essentials, 69 mid essentials, 77 late essentials, 493 neutral genes, and 100 non-targeting controls (NTCs) chosen randomly from the Yusa v3 NTCs. In the dual-guide benchmark library we additionally removed 8 neutral genes in which the mapping co-ordinates of several of the guides did not fall within the Ensembl gene co-ordinates (largely driven by mapping to paralogous genes) – since these guides were part of the original libraries they were not removed from the single-guide benchmark library to avoid any biases in the direct library comparisons.

### Benchmarking library cloning

A 75-mer oligo pool was ordered from Twist Bioscience. The oligo pool was PCR amplified using the KAPA HiFi HotStart ReadyMix (Roche #KK2602), 10ng of template, 300nM final concentration of primers oligo_pool1_for 5’-TATATATCTTGTGGAAAGGACGAAACACCG-3 and oligo_pool2_rev 5’-GCTGTTTCCAGCATAGCTCTTAAAC-3’. PCR cycling conditions: (1) 95°C for 3 min; (2) 98°C for 20sec; (3) 61°C for 15sec; (4) 72°C for 15sec; (5) 72°C for 1min; steps (2) to (4) were repeated for a total of 10 cycles. 3 independent PCR reactions were pooled and purified using the QIAquick PCR Purification Kit (Qiagen #28106).

The lentiviral gRNA expression vector pKLV3-U6gRNA5(BbsI)-PGKpuroBFP-W-L1 (Ong et al. 2017) was digested with BbsI-HF (NEB #R3539) and then gel purified. Amplicons were cloned into the digested vector using the NEBuilder HiFi DNA Assembly kit (#E2623S) according to manufacturer’s recommendation using a 5 to 1 molar ratio of amplicons to vector in three independent reactions. Gibson Assembly reactions were pooled, ethanol precipitated and resuspended in nuclease-free water. Three independent transformations of the Gibson Assembly reactions were carried out using NEB 10-beta electrocompetent E.coli (NEB #C3020K) according to manufacturer’s instructions. Bacteria were pooled and then cultured overnight at 30°C on two 225mm x 225mm LB Agar ampicillin plates with a coverage >500x per gRNA. Plasmid DNA was extracted using Macherey-Nagel NucleoSnap Plasmid Midi kit (#740494.50). NGS libraries from the plasmid DNA were prepared to confirm library representation and distribution.

### Full VBC top6 gRNA for benchmarking library

We added all missing gRNAs from the VBC top6 library (Vienna library) targeting our benchmarking gene list to our benchmarking library. For this purpose, an additional oligo pool containing 2155 gRNAs was ordered from Twist Bioscience and cloned together with the original benchmarking oligo pool for a total number of 25556 gRNAs into our newly generated pFGCLV1 backbone, starting from the PCR amplification as described above.

### Cloning of genome-wide Vienna single top3 and Vienna dual-guide libraries

The VBC top 6 sgRNA sequences (Homo sapiens hg38) were downloaded from: https://www.vbc-score.org/download. The top3 (auto-pick) sgRNAs/gene were selected (57307 gRNAs) and 500 non-targeting control guides (NTCs) from the Yusa v3 library were added for a total of 57807 gRNAs. Oligonucleotides were ordered from Twist Bioscience. PCR amplification, Gibson Assembly (GA) and electroporations were performed as described above with the exception that for the GA six independent reaction were performed followed by 8 independent electroporations. Bacteria were pooled and spread on 12 large LB Agar ampicillin plates and grown overnight at 30°C. Plasmid DNA was extracted using Macherey-Nagel NucleoBond Xtra Maxi kit (#740414.50). NGS libraries from the plasmid DNA were prepared to confirm library representation and distribution.

### Cloning of the genome-wide Vienna dual-guide and the dual-guide benchmarking libraries

Preparation of dual-guide libraries involved a two-step cloning process that was adopted from literature (Shen et al.; 2017). Briefly, oligo nucleotides were designed to contain pre-fixed combinations of two gRNAs targeting the same gene with a stuffer in the middle that contained two BsmBI restriction sites for the insertion of the second sgRNA scaffold and the mouse U6 promoter. Oligonucleotide synthesis and cloning of the 1^st^ step intermediate library was outsourced to Twist Bioscience. Subsequently, the 1^st^ step plasmid library, was digested by BsmBI and an insert containing a gRNA scaffold and the mouse U6 promoter was cloned in between the 1^st^ and 2^nd^ gRNA via ligation ensuring a library coverage >500-fold at all steps. NGS libraries from the 1^st^ and 2^nd^ step and the final plasmid library were prepared to confirm library representation and distribution.

For the genome-wide Vienna dual-guide library, we made combinations of guide RNAs using the VBC top1+3, VBCtop2+4 and VBCtop3+6 and added 250 control constructs paring 500 NTCs from the Yusa v3 library for a total of 57274 constructs.

For the dual-guide benchmarking library, we took the top4 gRNAs, based on the VBCscore, from the Brunello, Yusa v3, Croatan and Vienna library, and made pairs of 1+3, 2+4 and their reverse. Also, each guide RNA was paired with non-targeting guides in both directions to compare single vs. dual-targeting and the influence of the promoter and expression position on editing. The final dual-guide benchmarking library consisted of 45720 constructs. Oligo synthesis and cloning of the 1^st^ step dual-guide benchmarking library was also carried out by Twist Bioscience. Insertion of the first sgRNA scaffold and the mU6 promoter were performed in house. NGS libraries from the 1^st^ and 2^nd^ step and the final plasmid library were prepared to confirm library representation and distribution.

### Cell lines and culture

All cell lines were obtained from ATCC. Identities of all cell lines were confirmed via STR testing. All cell lines were also tested to be negative for mycoplasma prior to experiments. Cells were generally maintained in the absence of antibiotics except during screening (100U/ml penicillin and 100μg/ml streptomycin).

HCT116 and HT-29 were cultured in McCoy’s + 10% foetal bovine serum (FBS) + 1% GlutaMAX

SW480 and HEK293T were cultured in DMEM + 10% FBS + 1% GlutaMAX

RKO were cultured in MEM + 10% FBS + 1% GlutaMAX

A549 were cultured in Ham’s F-12K (Kaighn’s) Medium + 10% FBS + 1% GlutaMAX

HCC827 and PC-9 were cultured in RPMI 1640 + 10% FBS + 1% GlutaMAX

Cell lines stably expressing Cas9 were generated by transduction using pKLV2-EF1a-Cas9Bsd-W (Addgene #68343) lentiviral particles in the presence of 8μg/ml polybrene (Merck) followed by Blasticidin selection starting from day 2 post transduction. Cas9 editing efficiency in all cell lines was confirmed to be >90% based on tests using the fluorescent protein editing reporter pKLV2-U6gRNA5(gGFP)-PGKBFP2AGFP-W (Addgene #67980).

### Lentivirus production

20 million HEK293T cells were seeded into a T175 flask the day before transfection to reach ∼90% confluency the next day. On the day of transfection, 30μg of a transfer vector, 25μg of psPax2 (Addgene #12260) and 10μg of pMD2.G (Addgene #12259) were added to 3ml of Opti-MEM (ThermoFisher). Then, 195μl X-tremeGENE HP (Merck) was added, gently mixed and incubated for 20min at room temperature. The transfection complexes were carefully added to the HEK293T cell. Next morning, media was removed and 30ml fresh complete DMEM was added. ∼55h after transfection, supernatant containing lentiviral particles was harvested and filtered through a 0.45μm filter. Cleared viral supernatant was then aliquoted and stored at -80°C before use.

### Single and dual-gRNA benchmarking screens with RKO, HCT116, HT-29, SW480 and A549

20 million of cells were seeded into a triple layer flask together with an appropriate amount of benchmarking library virus in the presence of 8μg/ml polybrene (Merck). Transductions for all cell lines were performed in biological duplicates. Next day virus containing media was replaced with fresh media. Two days post transduction, percentage of BFP contained in the library vector was determined by flowcytometry. In all cases a transduction rate of ∼30% was reached which corresponded to at least a coverage of >200 cells per gRNA. Puromycin selection was initiated 2 days post transduction and maintained until the percentage of BFP^+^ cells were >90%. Cells were maintained to have a library coverage of at least 500 cells per gRNA. Cell pellets were collected on days 4, 7, 10, 14, 17, 21 and 25 post transduction and stored at -20°C before processing.

### Genomic DNA isolation and Next Generation Sequencing (NGS)

Genomic DNA (gDNA) was isolated using the QIAamp DNA Blood Maxi Kit (Qiagen) according to manufacturer’s instruction. NGS libraries were prepared in a two-step process. First, the integrated lentiviral cassette containing the gRNA was amplified from gDNA. For single gRNA libraries, primer NGS_v3_for 5’-ACACTCTTTCCCTACACGACGCTCTTCCGATCTCTTGTGGAAAGGACGAAACA-3’ and reverse primer NGS_v3_rev 5’-GTGACTGGAGTTCAGACGTGTGCTCTTCCGATCTACCCAGACTGCTCATCGTC -3’ was used. For the dual-guide samples the reverse primer NGS_dg_rev 5’-GTGACTGGAGTTCAGACGTGTGCTCTTCCGATCTTGCTATGCTGTTTCCAGCA-3’ together with the NGS_v3_for was used. PCR reactions with 5μg gDNA (single) or 2μg (dual-guide) per well were set up using the Q5 Hot Start High-Fidelity 2× Master Mix (NEB #M0494) in a total volume of 50μl. PCR reactions were scaled accordingly to amplify the gRNAs at a coverage of at least 200-fold. The PCR products were then pooled in each group and purified using QIAquick PCR Purification Kit (Qiagen). Final NGS libraries were generated using 2ng of the purified 1^st^ PCR product using the dual-indexing Illumina-compatible DNA HT Dual Index kit (Takara #R4000660,R400661). 2^nd^ PCR products were purified with AMPure XP beads (Beckman Coulter) at an 0.7 ratio. Purified 2^nd^ step libraries were quantified using the Qubit dsDNA Quantification Assay Kit (ThermoFisher) and sequenced on either HiSeq4000 by PE50bp or NovaSeq6000 PE100bp with a 30% PhiX spike-in.

### Genome-wide Osimertinib screens comparing Yusa v3, Vienna single top3 and Vienna dual-guide

HCC827-Cas9 (doubling time ∼44h) and PC-9-Cas9 (doubling time ∼29h) cell lines were transduced with the respective libraries in the presence of 8μg/ml polybrene to achieve a transduction rate of ∼30%. To maintain the same coverage of each library, for the Vienna single top3 and Vienna dual-guide, half the number of cells was transduced as these libraries contain roughly half the number of constructs. Two days post transduction, the percentage of BFP^+^ cells was checked by flowcytometry. Puromycin selection was started and maintained for 4 days until %BFP was >90%. At that point a baseline pellet was collected from all conditions. Cells were then split into the dimethyl sulfoxide (DMSO) and the treatment arms with two technical replicates in each. For HCC827 and PC-9, Osimertinib was used at a concentration of 10nM and 7.5nM, respectively, to achieve a growth inhibition of ∼80%, which was determined empirically in a previous experiment. During the screen, DMSO treated control cells were split every 3/4 days. Cell pellets were collected at every split, 7 for HCC827 and 5 for PC-9. For the Osimertinib treated cells due to the growth inhibition, media was changed at the same time as for the DMSO control, containing fresh compound up until day 16 (HCC827) or day 10 (PC-9) post baseline when the cell numbers increased enough to be split and to collect pellets. For HCC827 cells, pellets from the Osimertinib arm were collected on days 16, 19, 23 and 26 for the Yusa v3 and Vienna single top3 libraries, and days 14, 21 and 25 for the Vienna dual-guide. For the PC-9 Osimertinib arm, for all three libraries, cell pellets were collected on day 10 and 17.

NGS library preparation for the genome-wide screens was performed as described above for the benchmarking screens. Number of PCR reactions per sample and library was kept constant at 200-fold. Sequencing coverage was also kept constant and performed according to library size.

### Analysis

For each CRISPR pooled screen, DNA sequencing data was generated by either the Illumina HiSeq 4000 or NovaSeq 6000 platforms and FASTQ files were produced using bcl2fastq2. The reads were cross-referenced against the relevant gRNA library sequences and exact matches were counted to produce a gRNA count matrix (gRNAs as rows, samples as columns) using an AstraZeneca in-house CRISPR counting algorithm (unpublished). The resultant count matrices were then processed by CRISPRcleanR version 2.2.1 (22) and the log_2_ fold-change estimates relative to the plasmid (averaged across the two replicates) were used for plotting. Count matrices were also processed through Chronos (14) so that gene fitness estimates could be calculated across all time points simultaneously (a docker image and python scripts are available in the associated data and code -see the ‘Availability of data and materials’ section).

For the Osimertinib drug-gene interaction screen, treated and control sample count data were also processed through MAGeCK version 0.5.9.5 (20) and the gene-level log_2_ fold-changes (relative to the plasmid or contrasted between the treatment and control samples) and adjusted p-values for the treatment-vs-control contrasts were extracted and used to compare the three gRNA libraries. Precision-Recall curves were calculated using the control samples from the three screens and processing them through Chronos. The curves were calculated using the R package PRROC version 1.3.1 (23). In addition, a two-sample Chronos analysis was conducted by treating DMSO and Osimertinib-treated samples as separate cell lines in the Chronos analysis and contrasting their resultant gene fitness estimates to determine the treatment-versus-control effect size differential.

Gene expression data for the cell lines used in this study was downloaded from the Cancer Cell Line Encyclopedia (CCLE) project via the Broad DepMap portal (file: ‘OmicsExpressionProteinCodingGenesTPMLogp1.csv’). To compare log_2_ fold-changes for dual- and single-targeting guides, genes in the Neutral set were selected if they had a log_2_ TPM (transcripts per million) value of zero in the relevant cell line. Neutral genes with zero CCLE expression were used when calculating Precision-Recall curves.

All figures were plotted using R version 4.2.2 and the R packages ggplot2 version 3.4.2, gt version 0.10.0, ggpointdensity version 0.1.0, and UpSetR version 1.4.0 (24). A complete list of R package dependencies is provided in the figure code repository (see the ‘Availability of data and materials’ section).

## Author contributions

UM, GJH, DRT, AK, SL and DW conceived the project. SL and NG planned the experiments. SL, NG, AP, and KS performed the experiments. AK analysed the data. AK, NG, SL and DW wrote the manuscript. UM, GJH, DRT and DW supervised the project. All authors reviewed and approved the manuscript.

## Acknowledgements

We are very grateful to our colleagues Marica Gaspari, Lu Li, Andy Sayer, Curtis Hart, Josh Tweedy, Patryk Slazak, Keerthi Thelakkad Chathoth, Robin Stephenson, Diego Fontecilla and other members of the FGC team for their contributions and useful advice.

