## Supplementary Figures for "A benchmark comparison of CRISPRn guide-RNA design algorithms and generation of small single and dual-targeting libraries to boost screening efficiency"

a

b

c

d

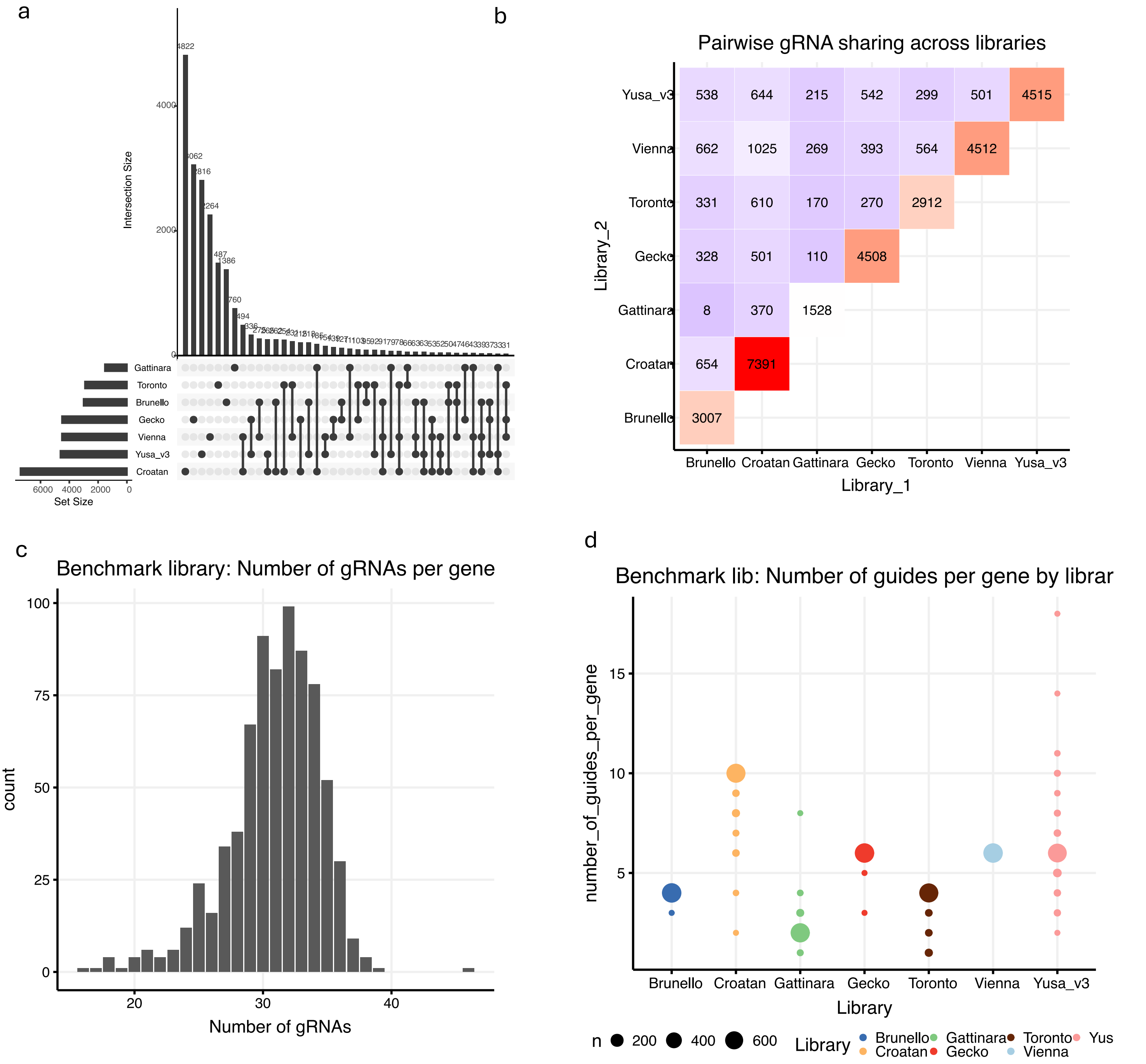


e

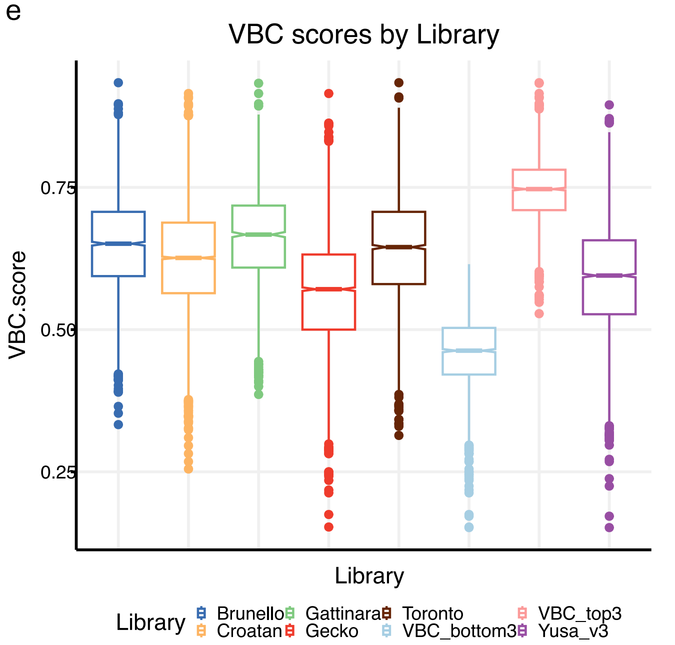


f

**Ontology enrichments**

$-\log_{10}$ (Adj. P-value)

| | Essential Genes | | |
|---|---|---|---|
| | Early | Mid | Late |
| **Spliceosome** | | | |
| KEGG SPLICEOSOME | 58 | | |
| RNA SPLICING | 33 | | |
| SPLICEOSOME ASSEMBLY | 9 | | |
| **Transcription** | | | |
| KEGG RNA POLYMERASE | 13 | | |
| TRANSCRIPTION INITIATION | 9 | | |
| TRANSCRIPTION | 5 | 4 | |
| **Ribosome** | | | |
| KEGG RIBOSOME | 49 | | |
| PROTEIN RNA COMPLEX ASSEMBLY | 19 | | |
| TRANSLATIONAL INITIATION | 8 | | |
| **Cell Cycle** | | | |
| KEGG CELL CYCLE | 14 | | |
| MITOSIS | 6 | 2 | |
| DNA REPLICATION | 5 | 3 | |
| **Metabolism** | | | |
| MITOCHONDRION | | | 18 |
| MITOCHONDRIAL RIBOSOME | | | 14 |
| KEGG RNA DEGRADATION | | | 3 |
| PHOSPHOLIPID BIOSYNTHETIC PROCESS | | | 3 |

**Figure S1. The distribution of gRNAs in the benchmark library.** a) An upset plot showing gRNA sharing across the 7 component libraries. b) A heatmap showing the number of guides shared by library pairs in the benchmark library. Since many of these pairwise shared guides will be shared by 3 or more libraries, the off-diagonal numbers can't simply be summed to arrive at the total number shared. The diagonal numbers represent the total number of guides in each library. c) A histogram showing the distribution of the number of gRNAs per gene in the benchmark library. d) The number of guides per gene broken down by library. e) VBC scores by library. f) Ontology enrichments for the early, mid, and late essential genes defined by Tzelepis et al. (2016) and which formed the basis of the gene selection for the benchmark library in the current study.

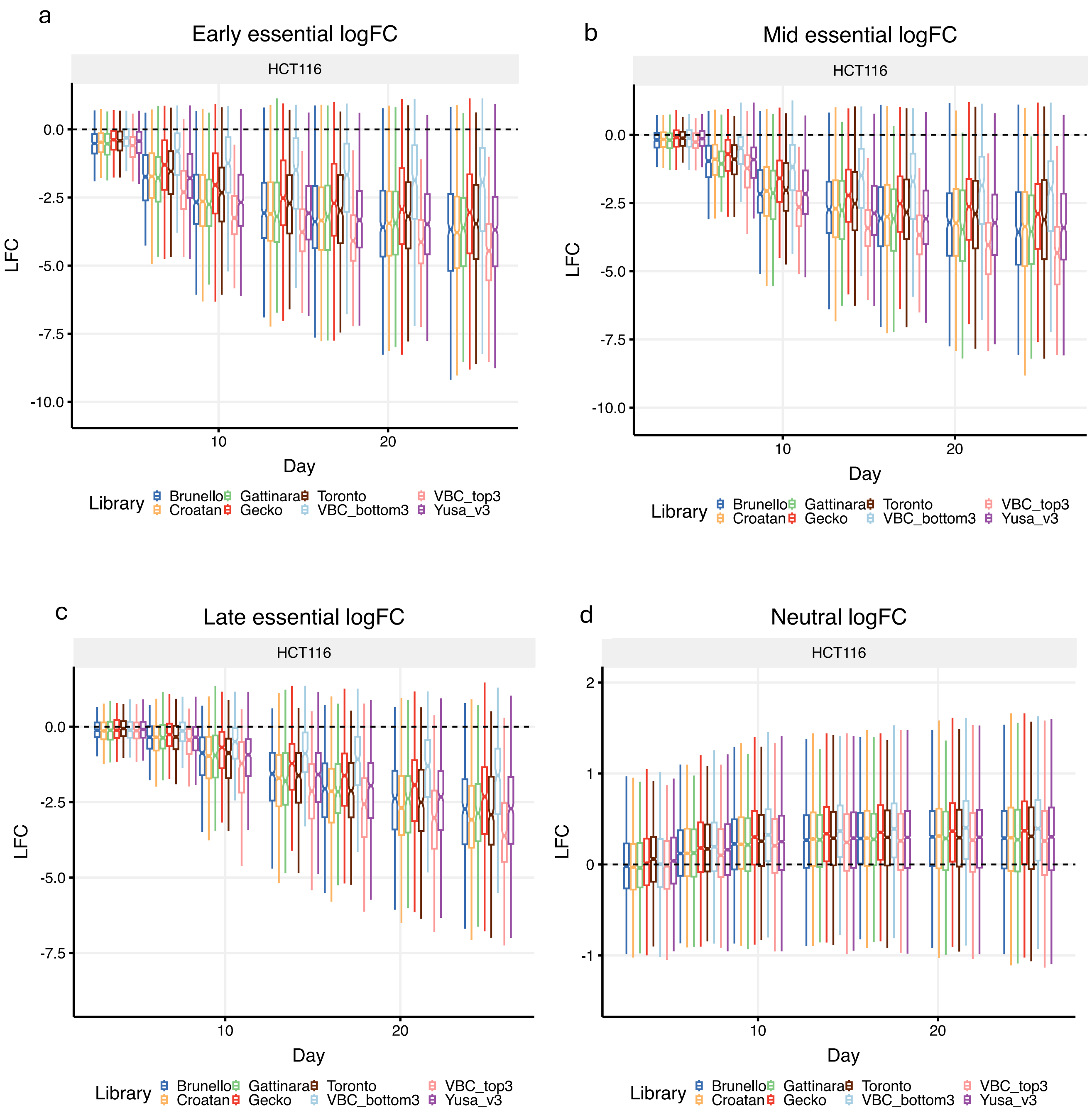

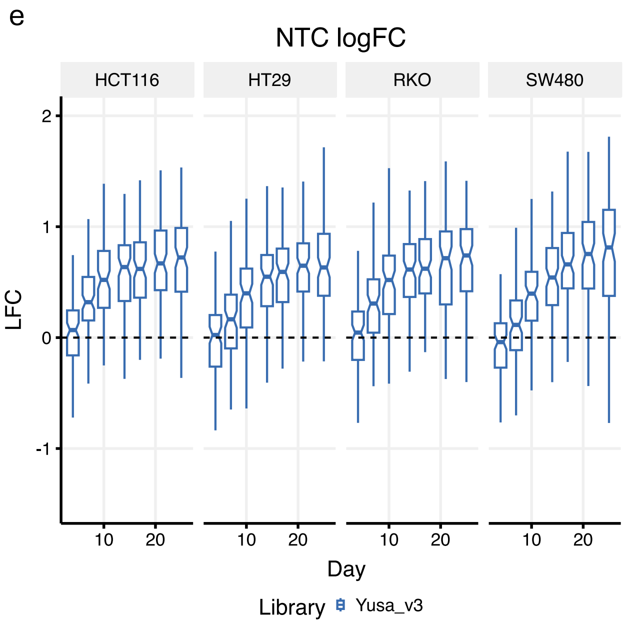


**Figure S2. Benchmark screen log-fold changes for different gene and guide classes.** a) Early essential depletion across 7 time points for HCT116. b) For mid essentials. c) For late essentials. d) Log-fold changes for neutral genes. e) Log-fold changes for 1000 Non-Targeting Control (NTC) guides from the Yusa_v3 library.

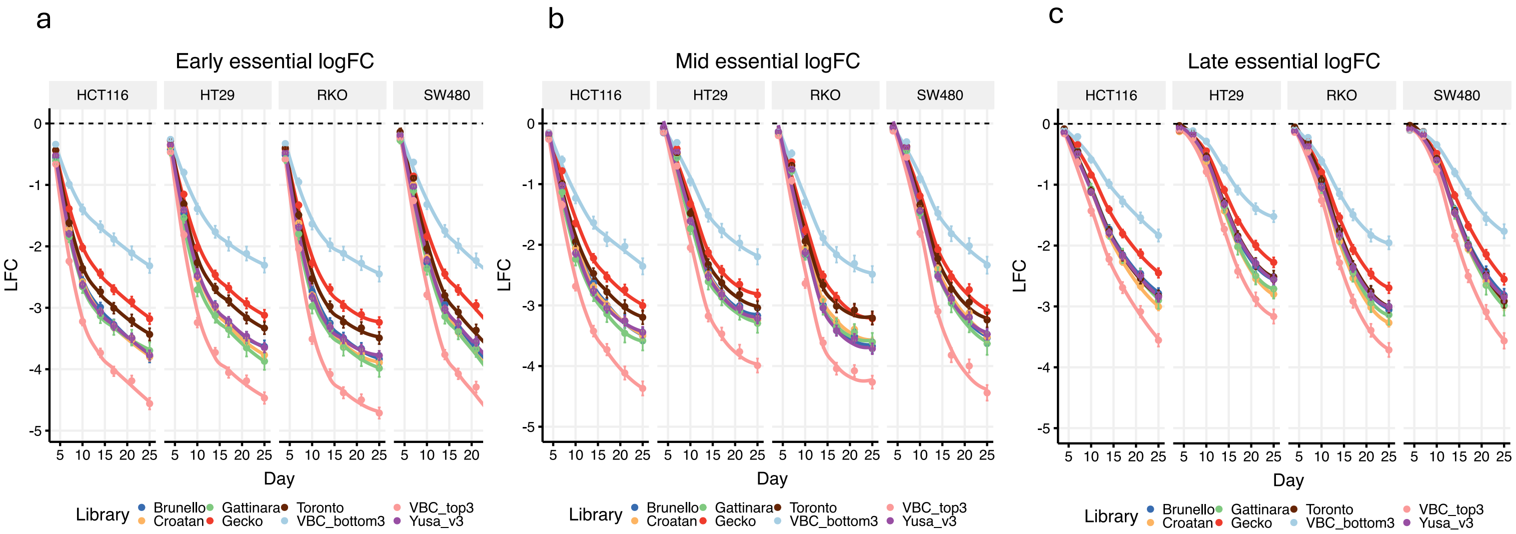


**Figure S3. Benchmark screen essential gene depletion.** a) Early essential log-fold changes across 7 time points and 4 cell lines. b) Mid essentials. c) Late essentials. Points mark average values across gRNAs and error bars are +/- the standard error of the mean. LOESS curves are fitted to the data.

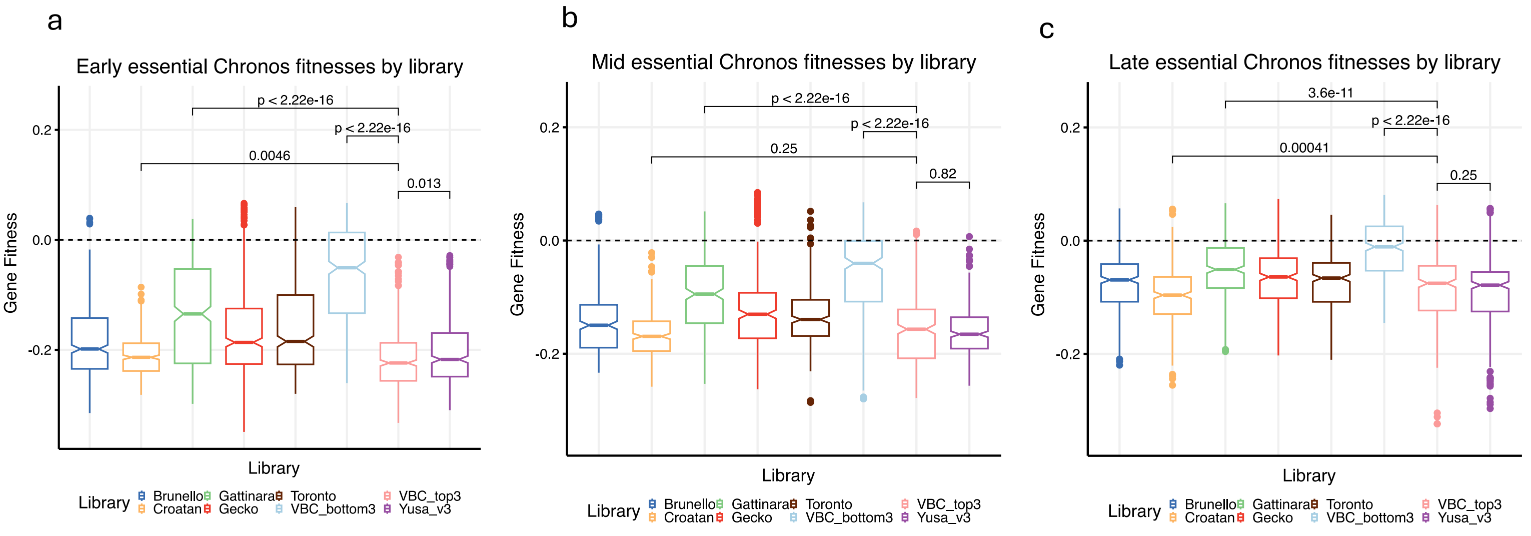


**Figure S4. Benchmark Chronos gene fitness estimates by library and pooled across 4 cell lines.** a) Early essential Chronos fitness estimates broken down by library. P-values are Wilcoxon two-sample tests. b) Mid essentials. c) Late essentials – the p-value between Croatan and VBC_top3 indicates Croatan is significantly more depleted at the gene level.

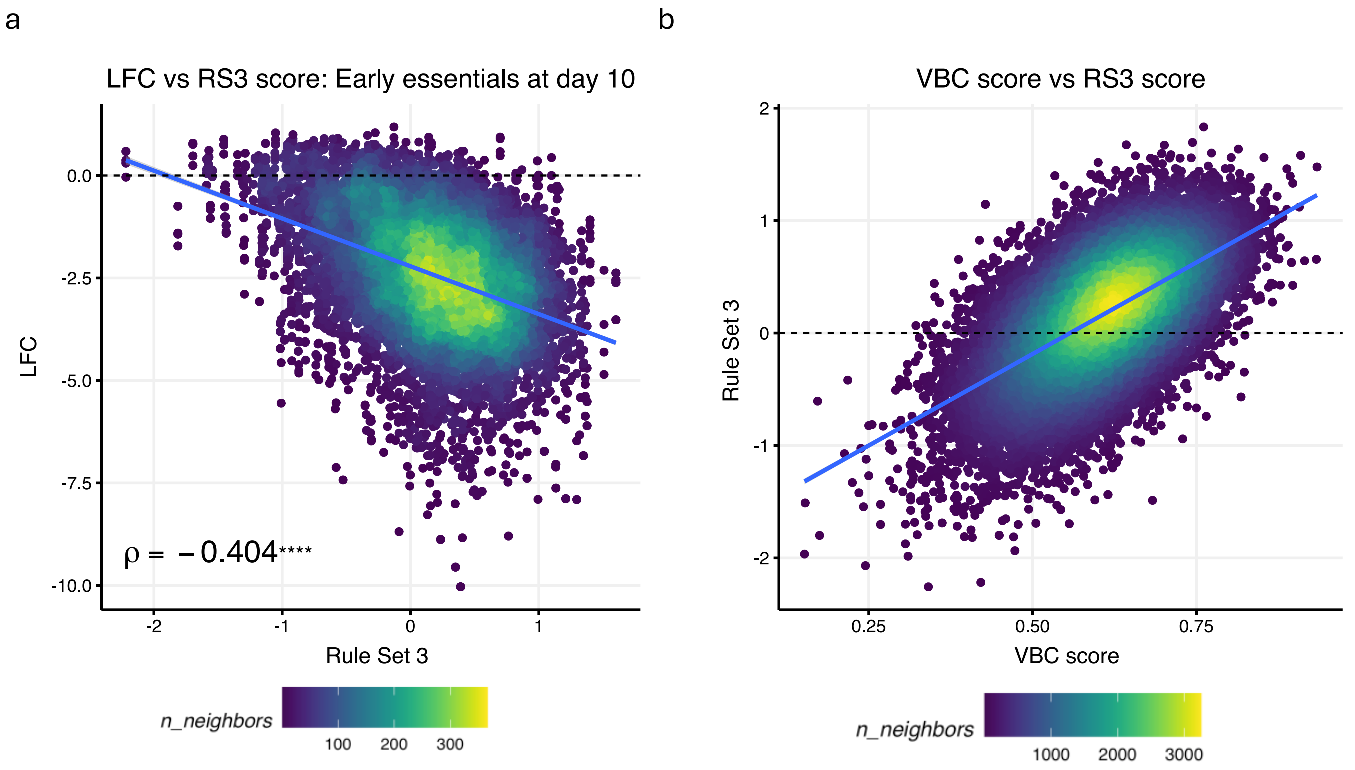


**Figure S5. Log-fold change correlation with Rule Set 3 scores.** a) Early essential depletion exhibits a significant negative correlation with Rule Set 3 scores. Spearman's Rho is shown. b) Correlation between VBC scores and Rule Set 3 scores.

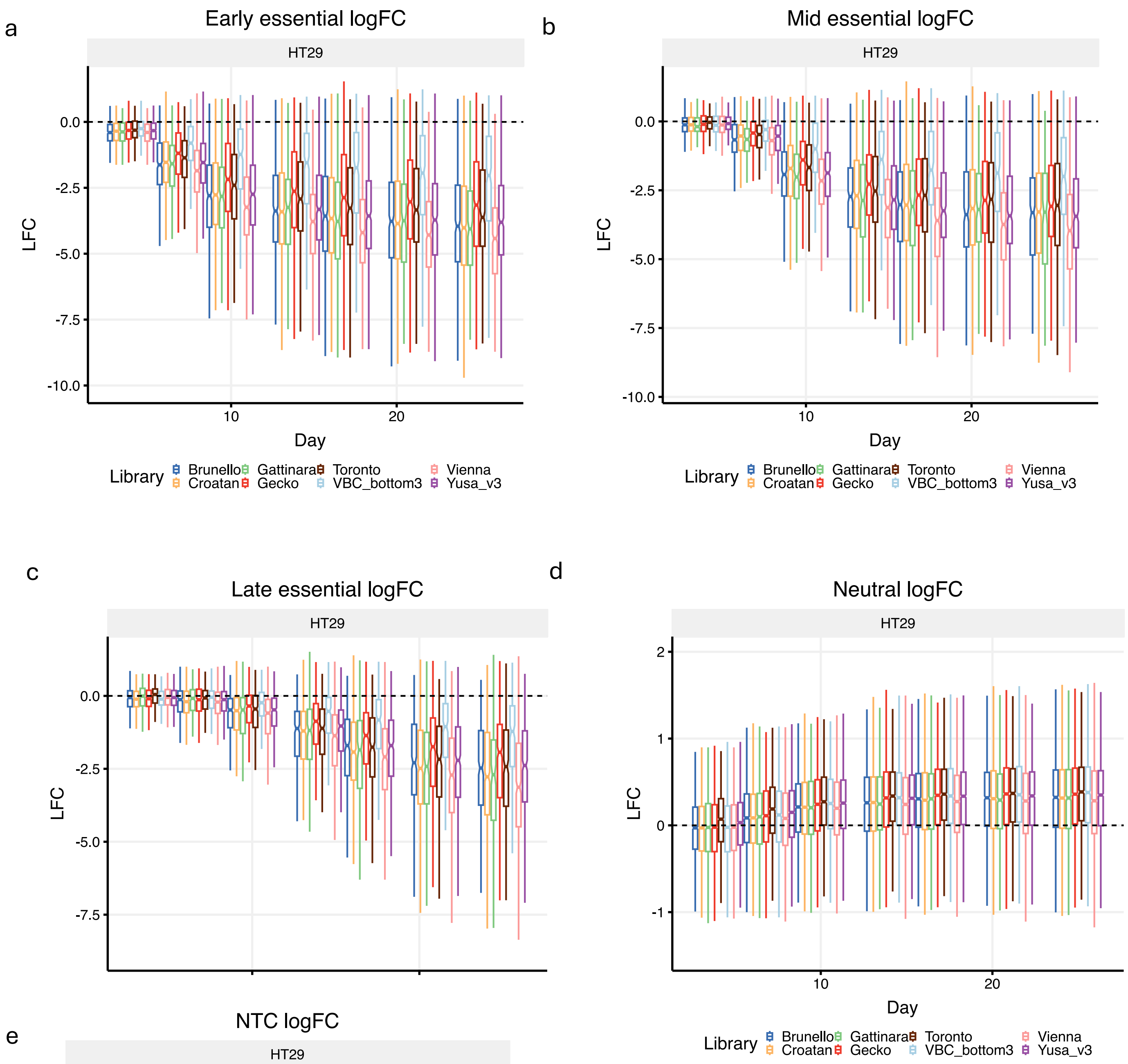


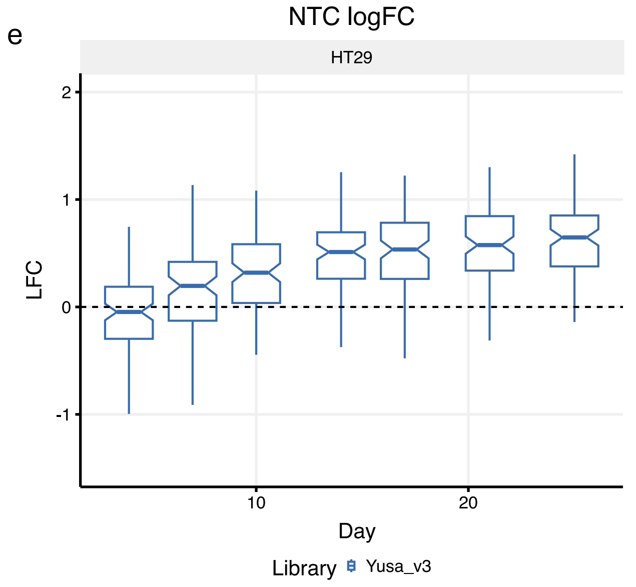


**Figure S6. HT29 benchmark screen log-fold changes for different gene sets – includes all-6 Vienna guides.** a) Early essential depletion across 7 time points for HT29. b) For mid essentials. c) For late essentials. d) Log-fold changes for neutral genes. e) Log-fold changes for 1000 Non-Targeting Control (NTC) guides from the Yusa_v3 library. For the ‘Neutral’ genes, the ‘VBC_bottom3’ guides are not identical to those shown in Fig. S2, they differ by 11 guides.

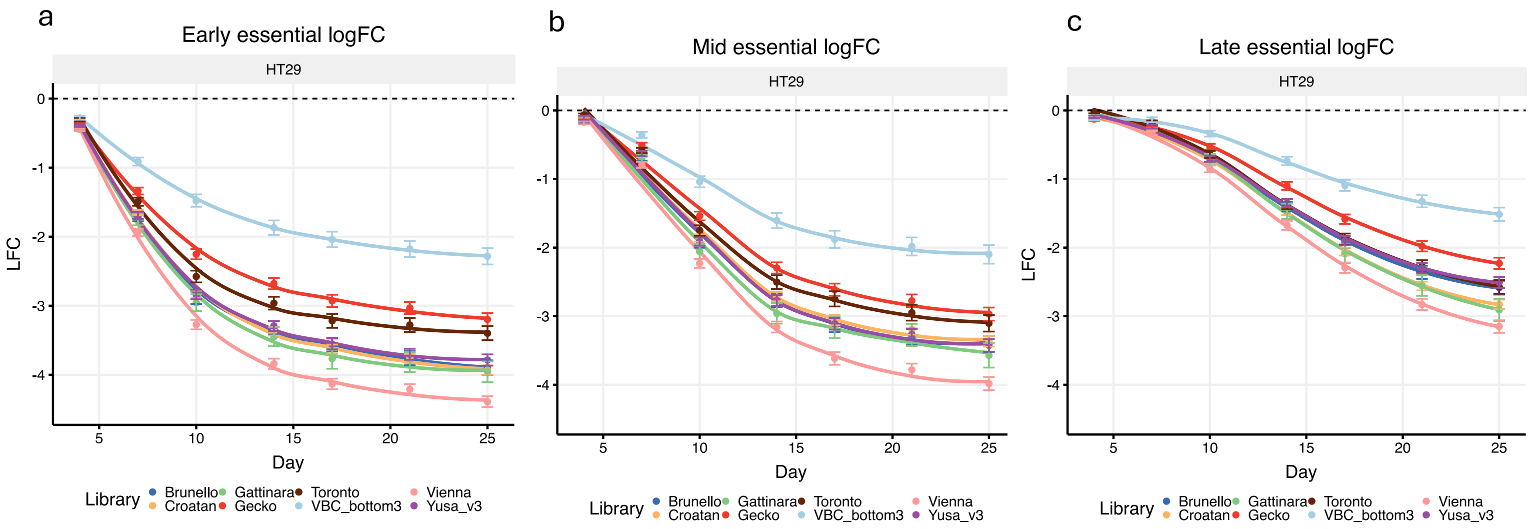


**Figure S7. Benchmark library essential gene depletion including the Vienna library.** a) Early essential log-fold changes across 7 time points and the HT29 cell line. b) Mid essentials. c) Late essentials. Points mark average values across gRNAs and error bars are +/- the standard error of the mean. LOESS curves are fitted to the data.

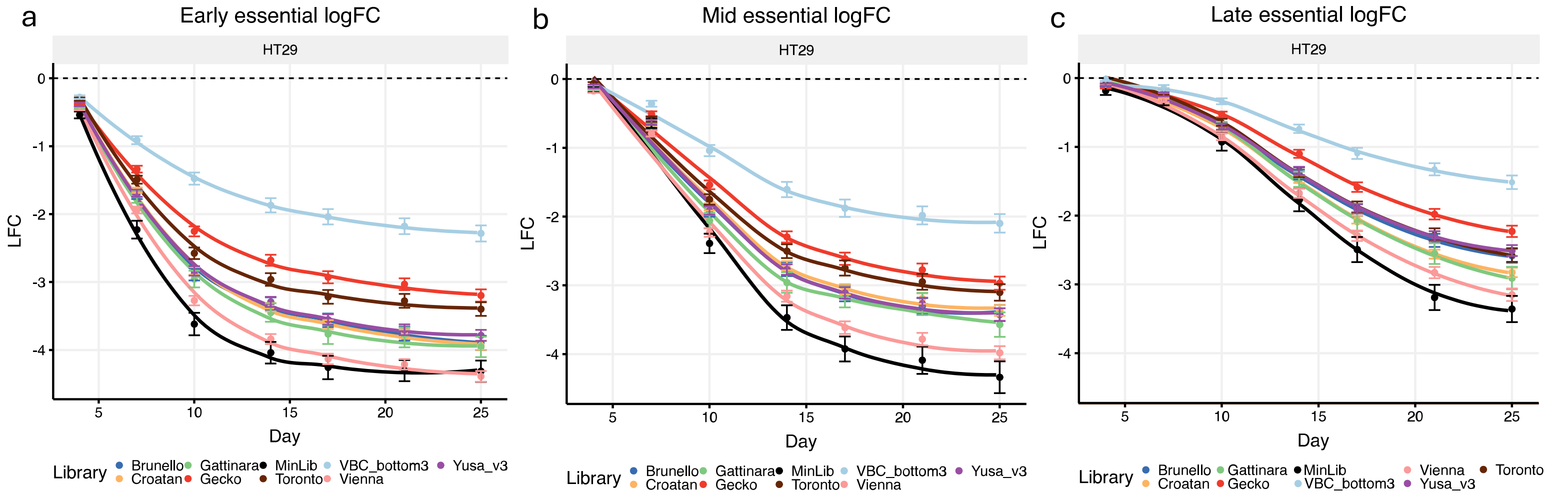


**Figure S8. Benchmark library essential gene depletion including 655 guides from the MinLib library.** a) Early essential log-fold changes across 7 time points and the HT29 cell line. b) Mid essentials. c) Late essentials. Points mark average values across gRNAs and error bars are +/- the standard error of the mean. LOESS curves are fitted to the data.

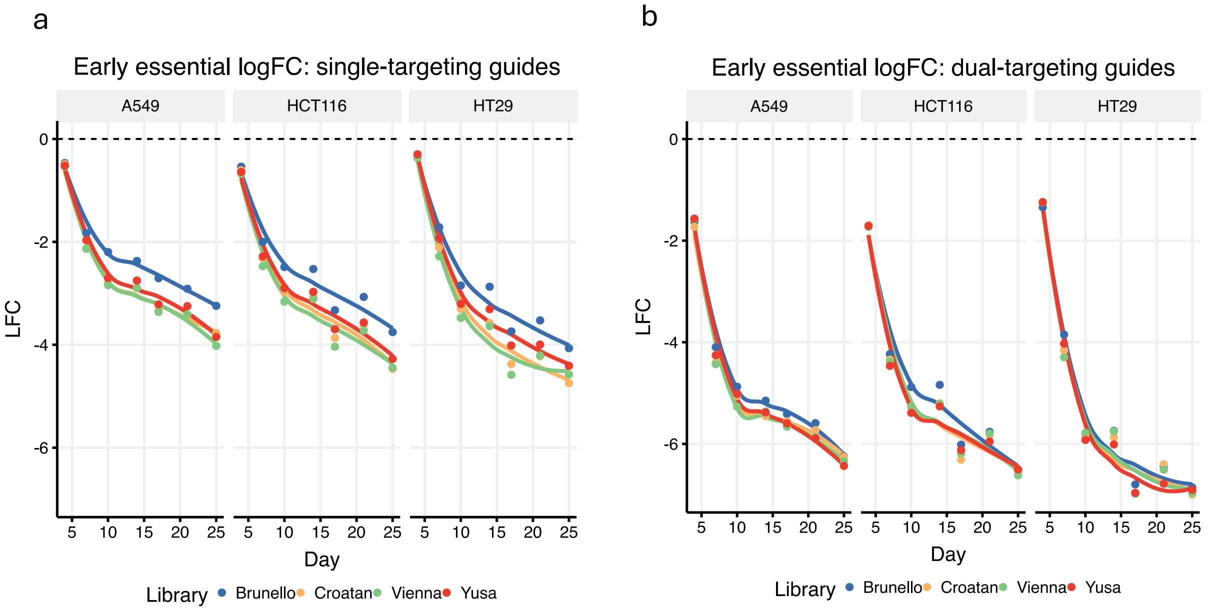


**Figure S9. Early essential depletion of single-targeting and dual-targeting guides broken down by component libraries.** a) Log-fold changes for single-targeting guides across 7 time points and 3 cell lines. b) Dual-targeting guides. Points mark average values across gRNAs and error bars are +/- the standard error of the mean. LOESS curves are fitted to the data.

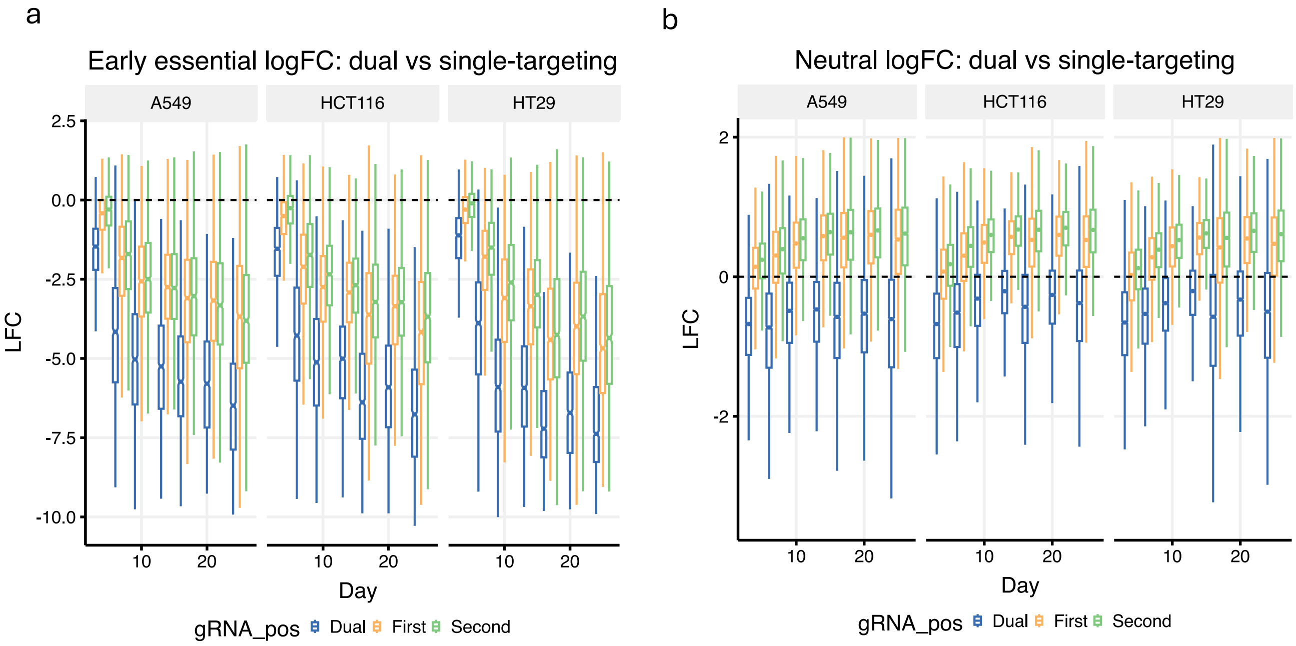


**Figure S10. Comparison of dual- and single-targeting guides for early essential and neutral genes.** a) Early essential depletion across 7 time points and 3 cell lines. b) For neutral genes. 'gRNA_pos' – gRNA position of the targeting guide, whether dual-targeting (Dual), or single-targeting in either the First or Second position.

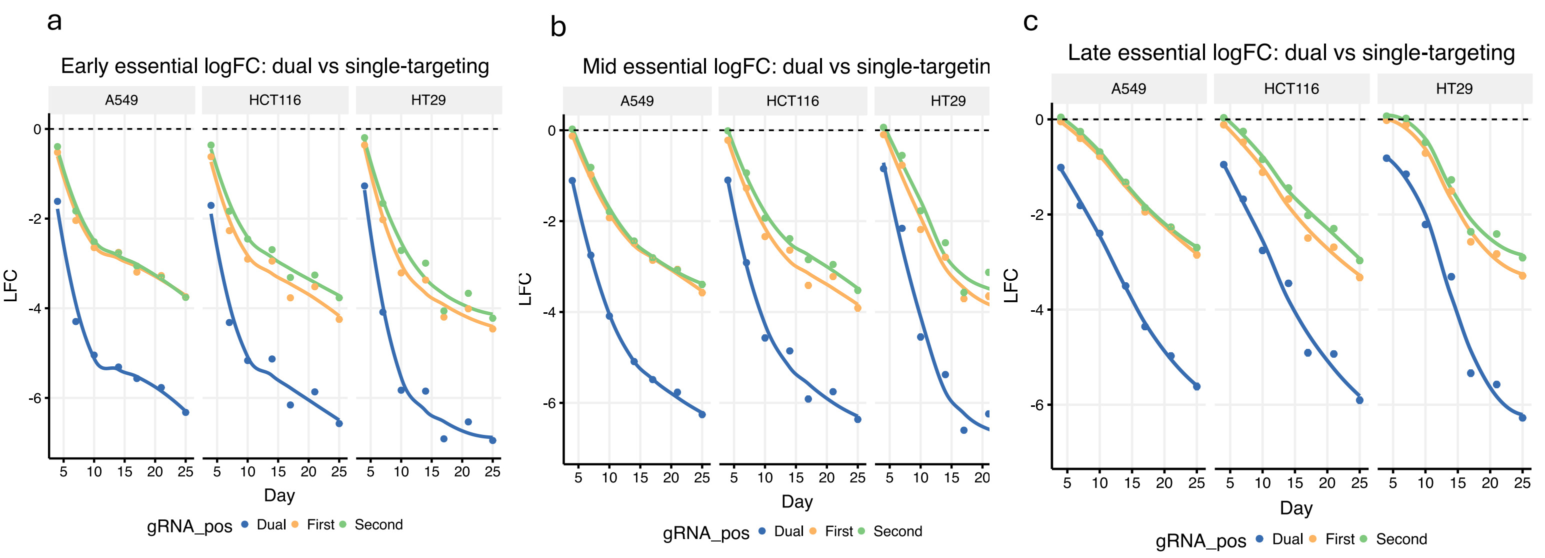


**Figure S11. Dual- and single-targeting essential gene depletion curves.** a) Early essential depletion across 7 time points and 3 cell lines. b) Mid essentials. c) Late essentials. ‘gRNA_pos’ – gRNA position of the targeting guide, whether dual-targeting (Dual), or single-targeting in either the First or Second position. Points mark average values across gRNAs and error bars are +/- the standard error of the mean. LOESS curves are fitted to the data.

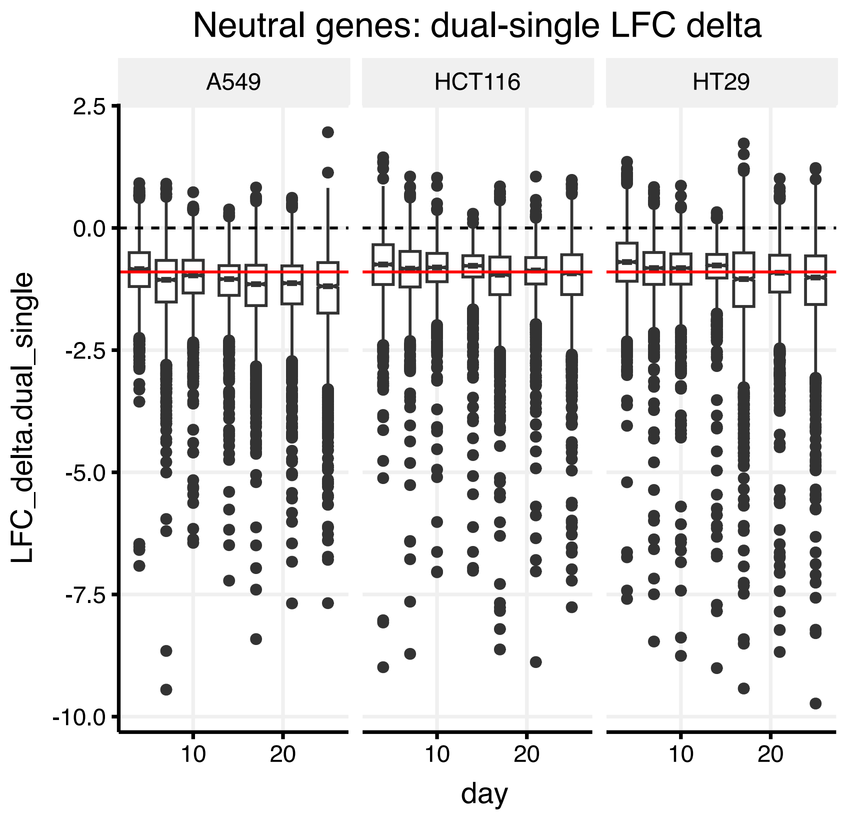


**Figure S12. Log-fold change deltas for dual-targeting vs-single-targeting gRNAs that target neutral genes with zero expression in CCLE data.** Log-fold change deltas as a function of time.

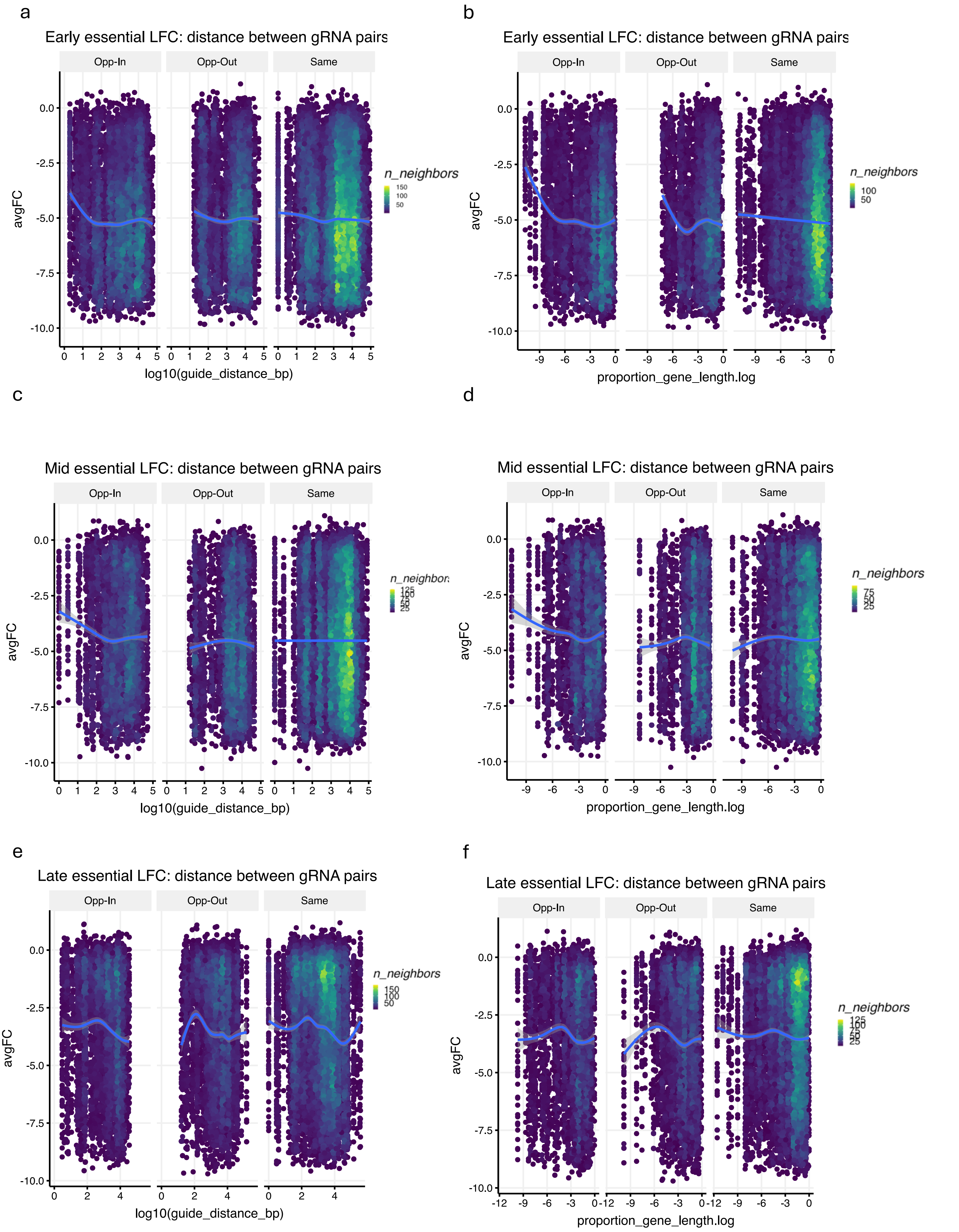


**Figure S13. Dual-targeting essential gene depletion as a function of gRNA pair distance.** a) Early essentials log10 absolute distance. b) Early essentials log proportion of total gene length. c) Mid essentials log10 absolute distance. d) Mid essentials log proportion of total gene length. e) Late essentials log10 absolute distance. f) Late essentials log proportion of total gene length. PAM pair orientation: Opp-In – opposite strands facing each other, Opp-Out – opposite strands facing away, Same – same strand.

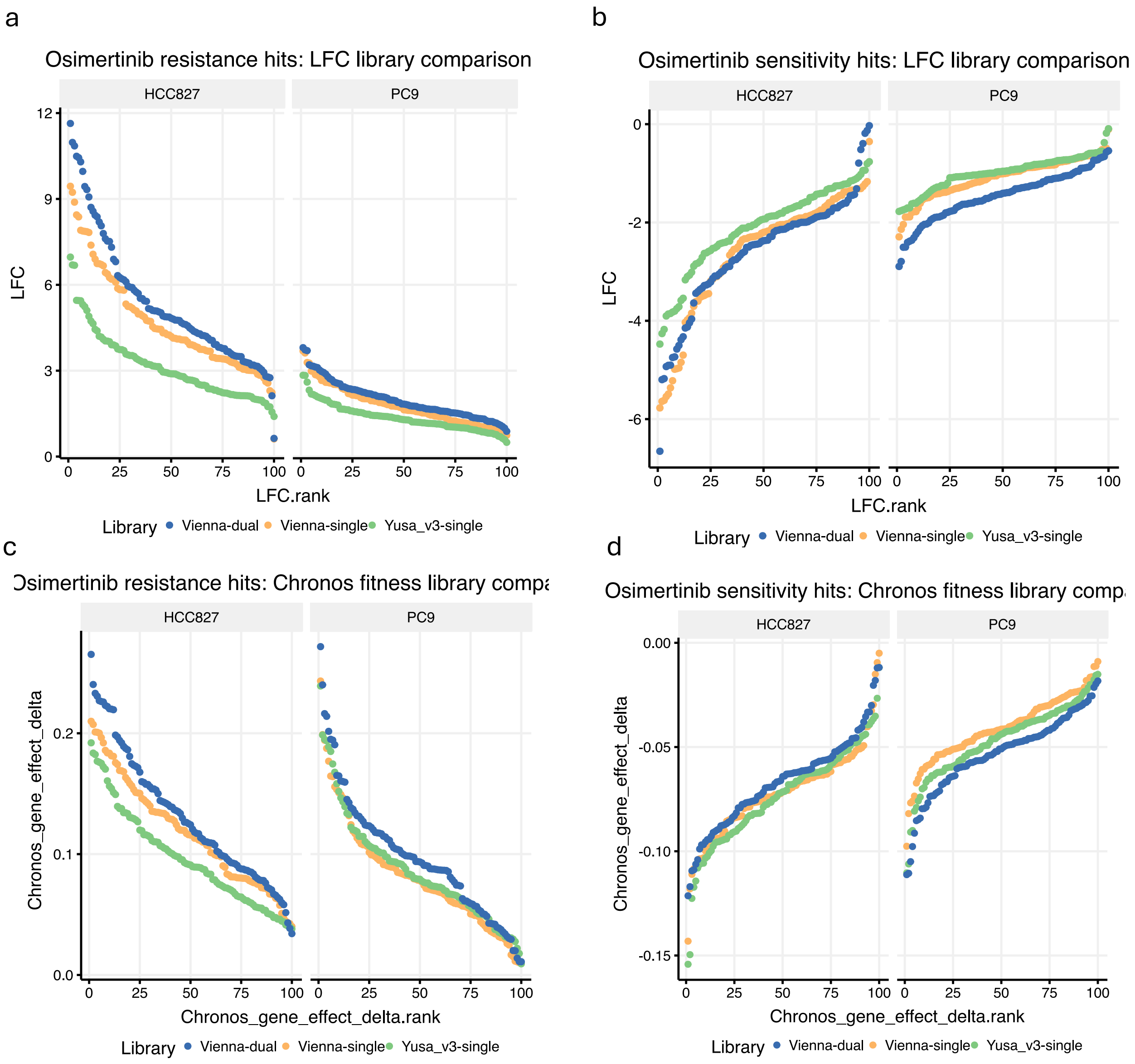


**Figure S14. Effect size library comparisons for the top 100 Mageck genes by FDR.** a) Resistance hit log-fold changes at approximately 10 doublings (day 19 for HCC827, day 10 for PC9). b) Log-fold changes for sensitivity hits. c) Resistance hit Chronos fitness estimates (estimated across all time points). d) Chronos fitnesses for sensitivity hits.

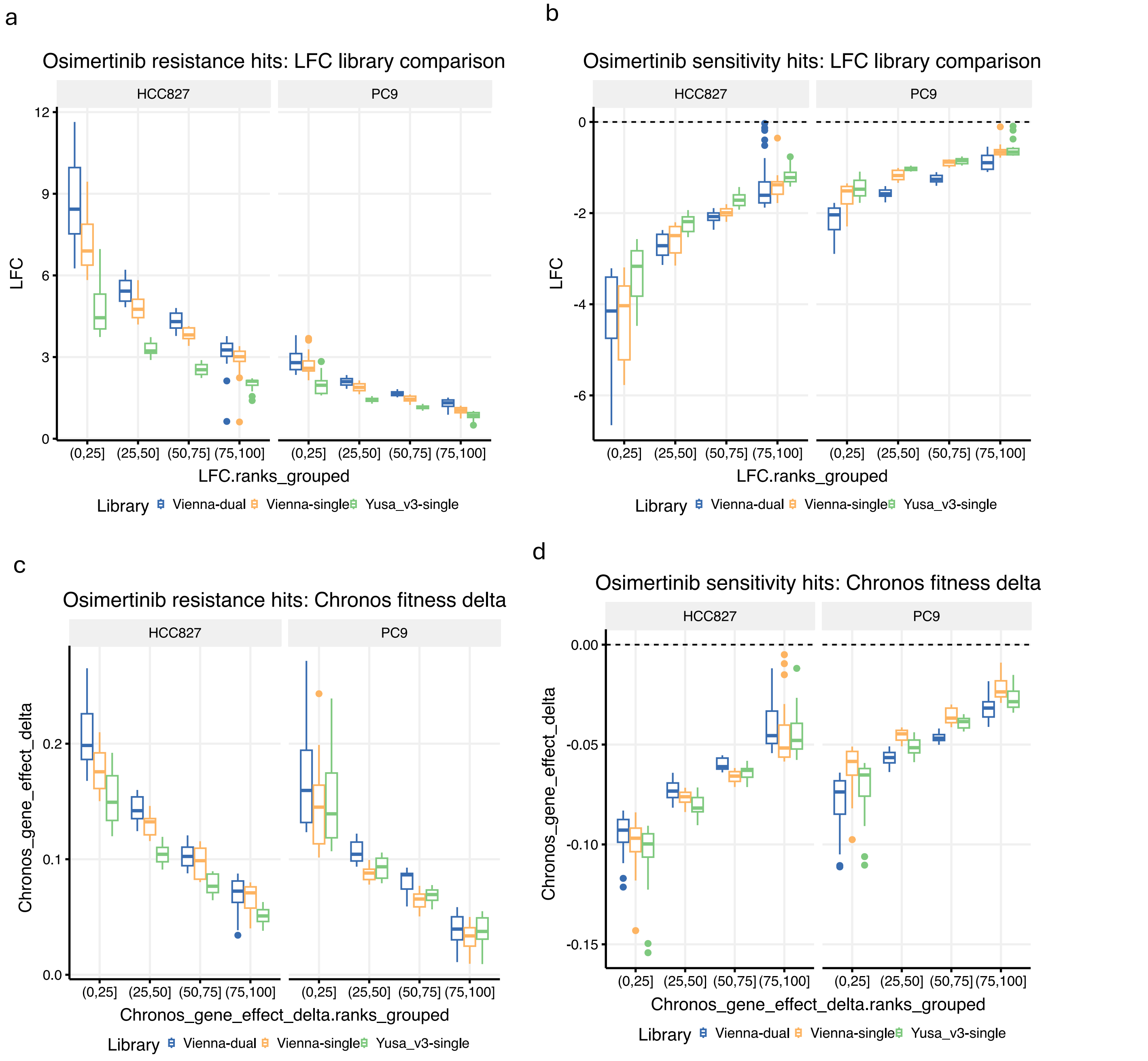


**Figure S15. Effect size library comparisons for top 100 Mageck genes by FDR grouped by ranks.** As in Figure S14 except ranked genes are grouped into bins. a) Resistance hit log-fold changes at approximately 10 doublings (day 19 for HCC827, day 10 for PC9). b) Log-fold changes for sensitivity hits. c) Resistance hit Chronos fitness estimates (estimated across all time points). d) Chronos fitnesses for sensitivity hits.
